# Comparative genomics reveals different population structures associated with host and geographic origin in antimicrobial-resistant *Salmonella enterica*

**DOI:** 10.1101/2020.01.28.923664

**Authors:** Jingqiu Liao, Renato Hohl Orsi, Laura Carroll, Martin Wiedmann

## Abstract

Genetic variation in a pathogen, including the causative agent of salmonellosis, *Salmonella enterica*, can occur as a result of eco-evolutionary forces triggered by dissimilarities of ecological niches. Here, we applied comparative genomics to study 90 antimicrobial resistant (AMR) *S. enterica* isolates from bovine and human hosts in New York state and Washington state to understand host- and geographic-associated population structure. Results revealed distinct presence/absence profiles of functional genes and pseudogenes (e.g., virulence genes) associated with bovine and human isolates. Notably, bovine isolates contained significantly more transposase genes but fewer transposase pseudogenes than human isolates, suggesting the occurrence of large-scale transposition in genomes of bovine and human isolates at different times. The high correlation between transposase genes and AMR genes, as well as plasmid replicons, highlights the potential role of horizontally transferred transposons in promoting adaptation to antibiotics. By contrast, a number of potentially geographic-associated single-nucleotide polymorphisms (SNPs), rather than geographic-associated genes, were identified. Interestingly, 38% of these SNPs were in genes annotated as cell surface protein-encoding genes, including some essential for antibiotic resistance and host colonization. Overall, different evolutionary forces and limited recent inter-population transmission appear to shape AMR *S. enterica* population structure in different hosts and geographic origins.

**Originality/Significance Statement:** *Salmonella enterica*, which is the causative agent of salmonellosis, poses a growing public health concern due to the emergence and spread of antimicrobial resistant (AMR) strains. The mechanisms underlying the population structure associated with different hosts and geographic origins of AMR *S. enterica* are underexplored due to limited genome-wide studies assessing the impact of ecological niches on genetic variations. By employing comparative genomics, our study provided insights into the genomic profiles of AMR *S. enterica* associated with two distinct hosts and two distant geographic locations, improving the mechanistic understanding of how bacterial population structure is shaped by different ecological niches. Our findings have broad implications for elucidating the impact of ecological and evolutionary forces on the adaptation, antimicrobial resistance, and pathogenicity of bacteria. Also, specific genetic markers we identified may help predict host or geographic origin of AMR *Salmonella* isolates, which could benefit the source tracking (e.g., host and geographic origins) of human disease cases and contamination events caused by AMR *S. enterica*.

## Introduction

Dissimilarity of ecological niches formed by different hosts or geographic origins is an important factor that can lead to genetic variation (e.g., gene presence/absence and polymorphism level divergence) and shape population structure, thus contributing to the evolution of pathogens (Strachan *et al.*, 2015; Hoberg and Brooks, 2008; Richards *et al.*, 2011; Strachan *et al.*, 2015). Besides providing insights in adaptation, pathogenesis and host range of pathogens, population structure associated with specific attributes (e.g., disease, host or geographic origin) also provides clues for evolutionary processes (e.g., selection, genetic drift and gene flow) that shaped pathogen diversity (Li *et al.*, 2019; Yue *et al.*, 2015). In addition, understanding of ecological niche-associated population structure is advancing the field of epidemiology by interrogating complex genome information to accurately predict contamination source and transmission patterns of pathogens including *Salmonella enterica* (Lupolova *et al.*, 2016; Lupolova *et al.*, 2017; Morgan *et al.*, 2004; Moura *et al.*, 2017).

The Gram-negative facultative anaerobe *S. enterica* is one of the major causes of human gastroenteritis, bacteraemia and enteric fever worldwide (Eng *et al.*, 2015; Ao *et al.*, 2015). The emergence and increasing prevalence of antimicrobial-resistant (AMR) *S. enterica* have further raised public health concerns and may increase the mortality rate of infections caused by this pathogen (Hong *et al.*, 2016; Eng *et al.*, 2015). AMR *S. enterica* has been detected in a wide range of animals in the majority of states in the US (Eng *et al.*, 2015; Centers for Disease Control and Prevention, 2017). Among >2,500 recognized serotypes of *S. enterica*, the broad host range serotypes Typhimurium and Newport are frequently associated with the human *Salmonella* infections, while the bovine adapted serotype Dublin is frequently associated with cattle infections. The emergence of AMR strains has been described for all three serotypes (Foley and Lynne, 2008). In our previous study of AMR *S.* Typhimurium, *S.* Newport and *S.* Dublin isolated from human and bovine sources in New York state (NY) and Washington state (WA), we found that it is likely that antimicrobial resistance has emerged independently in multiple *S.* Typhimurium lineages, as compared to single lineages of *S.* Newport and *S.* Dublin (Liao *et al.*, 2019). Also, our initial study of these isolates revealed a strong source and geographic association with antimicrobial resistance (Carroll *et al.*, 2017). For example, resistance to sulfamethoxazole-trimethoprim resistance was only observed in human isolates, while resistance to quinolones and fluoroquinolones was only observed in *S.* Typhimurium isolated from humans in WA. However, the mechanisms underlying the population structure associated with different hosts and geographic origins of AMR *S. enterica* are still underexplored at a genomic scale.

Comparative genomics is now widely used as a tool to study the evolution of bacteria (Chen *et al.*, 2006; Richards *et al.*, 2011; Zheng *et al.*, 2017) and can be used to identify genome-wide genetic variants that may be associated with host and geographic origin. Identification of such genetic variants in AMR *S. enterica* allows for a better understanding of genetic basis of pathogenicity and environmental adaptation. Humans and bovines are identified as two major hosts for AMR *S. enterica* (Dandekar *et al.*, 2015; Rodriguez-Rivera *et al.*, 2016; Zhao *et al.*, 2003). These two mammals differ in many aspects, e.g., physiology and diet, and hence form two distinct niches for associated microorganisms, including the pathogen *Salmonella*. While geographical origins can also represent distinct habitats and environmental conditions, definition of distinct niches based on geographical origins is more challenging since complex ecological factors (e.g., climate, anthropological activities) shape these niches. In order to assess the effects of both host and geographical origin on AMR *Salmonella* population structure, we selected to study isolates from two distant locations in the US: NY and WA. These two states showed similar incidence rate of reported *Salmonella* cases with 14.87 and 12.12 cases of salmonellosis per 100,000 population in NY and WA in 2015, respectively (Centers for Disease Control and Prevention, 2017). However, they represent distinct environments; NY has a humid continental climate, while the WA climate varies greatly from west to east with a rainy oceanic climate in the western half to an arid climate in the eastern half, where the majority of cattle operations are located. In addition, the management of dairy operations is different between NY and WA (United States Department of Agriculture, 2014). For example, the average dairy cattle herd size of WA is much larger (335 cows) than that of NY (113 cows). Consequently, NY and WA may represent distinct niches, particularly for bovine associated *Salmonella*. Hence, genomic comparison of AMR *Salmonella* isolated from bovine and human hosts in NY and WA will not only provide insights into the understanding of the adaption and evolution of this pathogen, but also has practical implications as genetic variation associated with different sources may allow for improved source tracking (e.g., human vs. bovine sources, different geographic sources) of human disease cases and contamination events. Thus, in this study, we used a previously reported WGS data set for 90 AMR *S.* Dublin, *S.* Newport, and *S.* Typhimurium isolates obtained from dairy cattle and humans from NY and WA (Carroll *et al.*, 2017; Liao *et al.*, 2019) to characterize population structure associated with host and geographic origin using comparative genomics.

## Materials and Methods

### Isolates and whole genome sequence data

A previously reported set of 90 *S. enterica* AMR isolates representing three serotypes including Dublin (n=21), Newport (n=32) and Typhimurium (n=37) collected between 2008 and 2012 (Carroll *et al.*, 2017; Liao *et al.*, 2019) was used for this study. Among those 90 isolates, 45 were collected from NY (Dublin (n=8), Newport (n=19) and Typhimurium (n=18)) and 45 were from WA (Dublin (n=13), Newport (n=13) and Typhimurium (n=19)). Also, among these 90 isolates, 41 were isolated from fecal samples of bovine (Dublin (n=10), Newport (n=14) and Typhimurium (n=17)) and 49 were isolated from the stool samples of human patients (Dublin (n=11), Newport (n=18) and Typhimurium (n=20)).

Genome sequence data for all 90 isolates have previously been reported (Carroll *et al.*, 2017) and deposited in the National Center for Biotechnology Information’s (NCBI) Sequence Read Archive (SRA) under accession number SRP068320. Genome assembly has also been reported previously (Carroll *et al.*, 2017, Liao *et al.*, 2019) and assembled genomes are available at NCBI DDBJ/ENA/GenBank under the accession numbers listed in Additional file 1: Table S4 in Liao *et al.*, 2019. Previously reported data (Liao *et al.*, 2019) for (i) genome annotation including identification of pseudogenes by NCBI Prokaryotic Genome Annotation Pipeline (Tatusova *et al.*, 2016), (ii) identification of orthologous genes (non-pseudogenes) by OrthoMCL (Li *et al.*, 2003), (iii) assignment of gene ontology (GO) and enzyme commission (EC) terms by Blast2GO v1.2.1 (Conesa *et al.* 2005), and (iv) high-quality core SNPs for *S.* Dublin, *S.* Newport, and *S.* Typhimurium called by Cortex variant caller (Iqbal *et al.*, 2012) were used here.

### Statistics

The Mann-Whitney test was carried out in Prism 7 to determine if the numbers of all pseudogenes as well as pseudogenes and functional genes annotated as encoding transposase differed significantly between bovine isolates and human isolates and if the numbers of all pseudogenes differed significantly between isolates from NY and WA as well. The false-discovery rate procedure (FDR) of Benjamini and Hochberg (BH) (1995) was employed in R version 3.6.0 to correct for multiple testing among isolate groups.

Non-metric multidimensional scaling (NMDS) (Kruskal, 1964) was performed using Bray-Curtis dissimilarities and the metaMDS function in R version 3.6.0’s vegan package to compare the dissimilarity of gene presence/absence pattern among isolates across serotypes and within each serotype. Permutational multivariate analysis of variance (PERMANOVA) (Anderson and Walsh, 2013) was employed using the adonis function in R version 3.6.0’s vegan package to test whether the centroids of isolate groups as defined by host or geographic origin are equivalent for all groups based on gene presence/absence among isolates across serotypes and within each serotype. *P*-values and PERMANOVA test statistics (F) were obtained using Bray-Curtis dissimilarities and 999 permutations. The FDR of BH was employed in R version 3.6.0 to correct for multiple testing among isolate groups.

Enrichment analyses of pseudogene functions, orthologous genes, and GO terms were conducted using Fisher’s exact test in R version 3.3.1 for (i) bovine versus human isolates and (ii) isolates from NY versus isolates from WA. FDR of BH was employed using R version 3.6.0 to correct for multiple testing for Fisher’s exact tests for pseudogene functions, orthologous genes, and GO terms. Items were defined as significantly enriched in one host or geographic origin when FDR was < 0.05 and odds ratio was > 6.71. These cut-offs were based on a previous report that odds ratio = 6.71 is equivalent to Cohen’s d = 0.8 (large), indicating strong association (Chen *et al.*, 2010).

Correlations between presence/absence of orthologous genes annotated as encoding transposase in this study and AMR genes as well as plasmid replicons previously reported by Carroll *et al.* (2017), using the same dataset, were assessed by Phi coefficient using the Python sklearn.metrics module. Transposase genes, AMR genes, and plasmid replicons that are present in all 90 genomes and less than 5 genomes were excluded from the Phi coefficient analyses. Items with Phi coefficient r > 0.25 were defined as having a strong positive correlation and included in heatmap visualization using R version 3.6.0.

Custom Python scripts were used to identify SNPs in (i) core genes present in all three serotypes and (ii) core genes defined separately for each of the serotypes. Fisher’s exact tests were employed to identify SNPs at which the distribution of nucleotides is dependent on host and/or geographic origin of isolates; these tests were performed in R version 3.3.1 followed by FDR of BH. Potentially geographic- or host-associated SNPs were defined as significant when FDR was < 0.05.

## Results

### Orthologous gene and pseudogene profiles differed significantly between bovine and human AMR *Salmonella* isolates

Overall, the genomes of the 90 AMR *S. enterica* isolates included 3,637 core genes (i.e., genes present in all 90 isolates) as well as 3,440 accessory genes. NMDS plots showed that across serotypes (Figure 1a) and within each serotype (Figure 1b, 1c, 1d), isolates clustered according to their host based on gene presence/absence, with no evidence for clustering by geographic location. PERMANOVA further showed that bovine and human cluster centroids, based on gene presence/absence, were not equivalent across serotypes and within each serotype (FDR < 0.01; Table 1), while no significant difference was observed between isolates from NY and WA in all tests (FDR > 0.05; Table 1).

**Figure 1.**
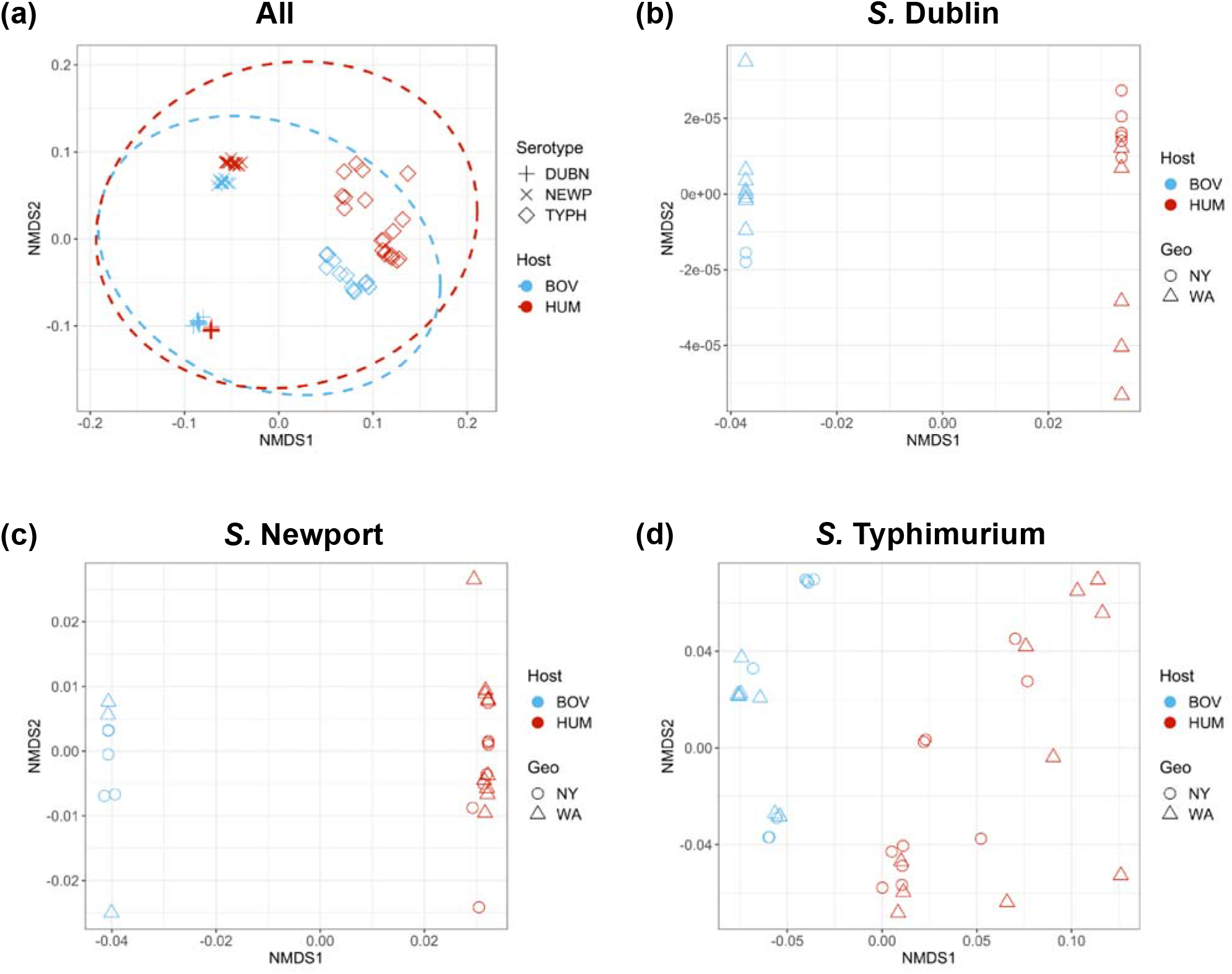
Gene presence/absence based-Non-metric multidimensional scaling ordination (NMDS) of antimicrobial-resistant (AMR) *S. enterica* isolates (a) across all serotypes; (b) within *S.* Dublin, (c) within *S.* Newport, and (d) within *S.* Typhimurium. Isolates from New York state (NY) are indicated by symbol circle; isolates from Washington state (WA) are indicated by symbol triangle. Bovine isolates (BOV) are in blue; human isolates (HUM) are in red. For panel (a), *S.* Dublin are indicated by “+”; *S.* Newport are indicated by “×”; *S.* Typhimurium are indicated by “◊”. Dotted lines indicate clustering of isolates correspond to host.

**Table 1.** PERMANOVA of gene presence/absence for isolate groups of AMR *S. enterica*^a^

| Serotypes | Grouping factor<sup>b</sup> | F Statistic | Uncorrected *p*-value | FDR<sup>c</sup> |
| --- | --- | --- | --- | --- |
| **All** | **Host** | **13.98** | **< 0.001** | **< 0.001** |
| All | Geographic origin | 0.86 | 0.48 | 0.48 |
| **S. Dublin** | **Host** | **21.37** | **< 0.001** | **< 0.001** |
| S. Dublin | Geographic origin | 2.75 | 0.04 | 0.06 |
| **S. Newport** | **Host** | **17.686** | **< 0.001** | **< 0.001** |
| S. Newport | Geographic origin | 2.24 | 0.05 | 0.06 |
| **S. Typhimurium** | **Host** | **10.37** | **< 0.001** | **< 0.001** |
| S. Typhimurium | Geographic origin | 1.21 | 0.26 | 0.29 |

<sup>a</sup> Rows in boldface indicate that FDR test was significant (FDR < 0.05).
<sup>b</sup> Host – bovine vs human, Geographic origin – NY vs WA.
<sup>c</sup> FDR was determined by a Benjamini and Hochberg correction.

The number of pseudogenes among bovine AMR *S. enterica* isolates were significantly lower than human isolates (Figure 2a*, P* < 0.0001). Across all serotypes, isolates from bovine and human sources had an average of 107 and 160 pseudogenes, respectively. Similarly, for each serotype bovine isolates had lower average numbers of pseudogenes; bovine isolates had 154, 89, and 94 pseudogenes, while human isolates had 225, 134 and 148 pseudogenes on average within *S.* Dublin, *S.* Newport and *S.* Typhimurium, respectively. By contrast, for all serotypes and within each serotype, the number of pseudogenes was not significantly different among isolates from NY and WA (Figure 2b); the AMR *S. enterica* isolates from NY and WA had an average of 132 and 140 pseudogenes, respectively. Within *S.* Dublin, *S.* Newport and *S.* Typhimurium, isolates from NY had an average number of 207, 109, and 123 pseudogenes, while isolates from WA had 181, 122 and 124 pseudogenes on average, respectively.

**Figure 2.**
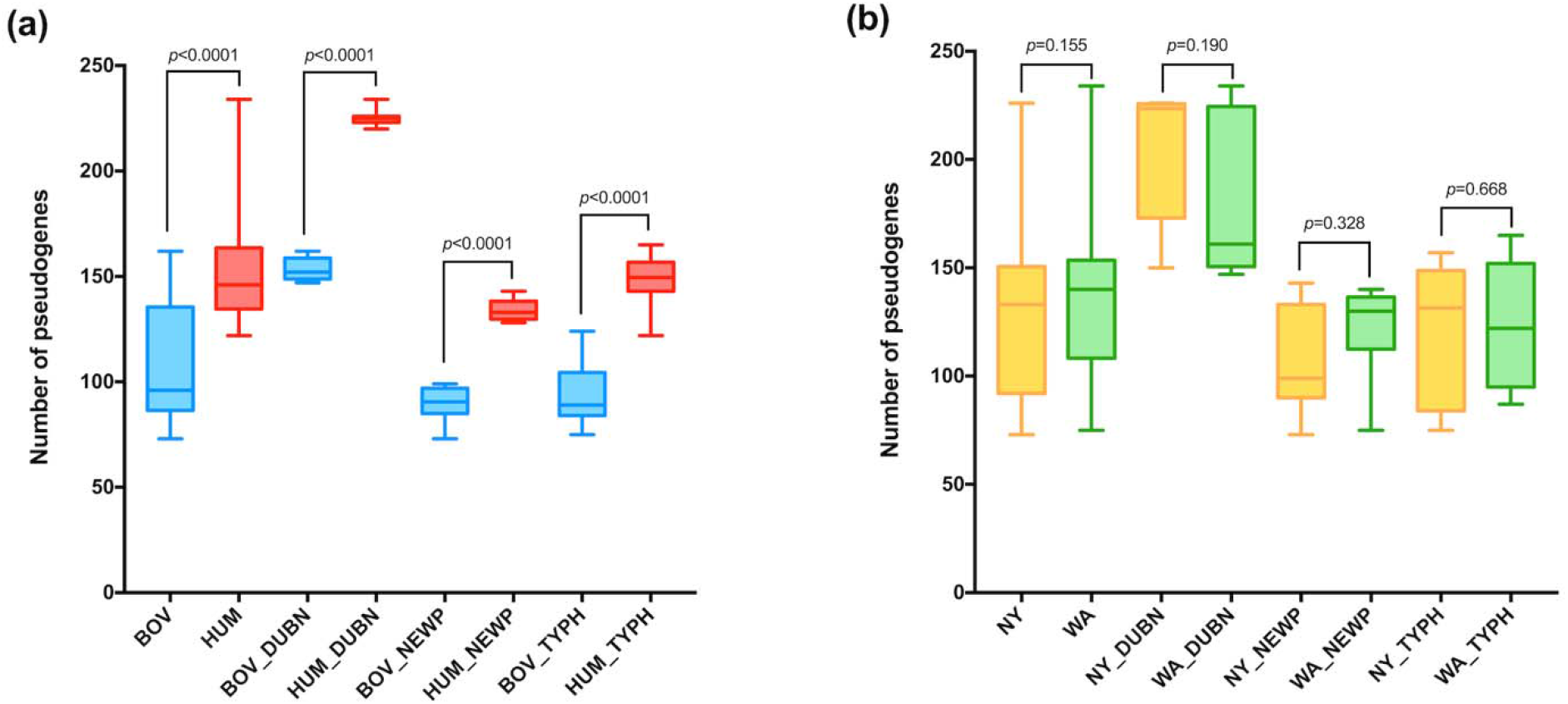
Box and whisker plot of the number of pseudogenes in AMR *S. enterica* isolates from (a) bovine and human populations, and (b) NY and WA among all three serotypes and within each serotype. DUBN - *S.* Dublin; NEWP - *S.* Newport; TYPH - *S.* Typhimurium; *P* values were determined by Mann-Whitney test after FDR correction. Minimum and maximum values are depicted by short horizontal lines above and below the box; the box signifies the upper and lower quartiles, and the median is represented by a short line within the box.

Enrichment analysis identified 55, 35, 33, and 34 pseudogene functions significantly enriched in human isolates across all serotypes, within *S.* Dublin, *S.* Newport, and *S.* Typhimurium, respectively (FDR < 0.05, odds ratio > 6.71; Table S1). Among the human-associated pseudogene functions, 8 (e.g., IS3, IS4, IS5/IS1182, IS630 family transposase), 27 (e.g., MarR family transcriptional regulator, effector protein YopJ), 20 (e.g., type VI secretion lipoprotein/VasD), and 14 (e.g., IS66 family transposase) were exclusively present in all human isolates of all serotypes, *S.* Dublin, *S.* Newport, and *S.* Typhimurium, respectively (Table S1). By comparison, enrichment analysis identified fewer pseudogene functions significantly overrepresented among bovine isolates across all serotypes (35 functions), within *S.* Dublin (20 functions), *S.* Newport (20 functions), and *S.* Typhimurium (23 functions) (FDR < 0.05, odds ratio > 6.71; Table S1). Among the bovine-associated pseudogene functions, 13 (e.g., arginine:ornithine antiporter, multidrug transporter subunit MdtG), 9 (e.g., acyl carrier protein), and 4 (e.g., PTS fructose transporter subunit EIIBC) were exclusively present in all bovine isolates representing Dublin, Newport, and Typhimurium, respectively (Table S1). By contrast, no significant geographic-associated pseudogene functions were identified except for two (alpha-ketoglutarate transporter and peptide transporter) found enriched in *S.* Newport isolates from NY.

### A number of genes were associated with bovine hosts, including 30 genes found exclusively in all bovine isolates

Among the 3,440 accessory genes for all 90 isolates, a total of 118 genes were significantly enriched in bovine isolates (FDR < 0.05, odds ratio > 6.71), including 30 genes found among all 41 bovine isolates regardless of serotype, but not present in any human isolates (Figure S1a; see Table S2 for a list of all genes). Annotated functions associated with these 30 genes predominantly represented the category “hypothetical proteins” (25 of the 30 genes); the remaining 5 genes were annotated as encoding transposase, small toxic polypeptide LdrA/LdrC, aspartate ammonia-lyase, permidine/putrescine ABC transporter substrate-binding protein, and racemase (Table S2).

Within *S.* Dublin, *S.* Newport, and *S.* Typhimurium, a total of 79, 64 and 82 genes, respectively, were significantly enriched in bovine isolates (FDR < 0.05, odds ratio > 6.71); these genes included 72, 54, and 46 genes exclusively found in all bovine isolates classified into a given serotype, but not identified in any of the human isolates classified into the same serotype (Figure S1a, Table S2). Based on these data we identified an overall number of 143 bovine-associated genes, which included the 118 genes identified as bovine-associated across all serotypes as well as additional bovine-associated genes found within each serotype. Among all these 143 bovine-associated genes, 60 genes were annotated with known functions; 19 and 11 of these 60 genes were annotated as encoding transposases and cell surface proteins, respectively (Figure 3, Table S3). The annotated functions for 15 bovine-associated genes matched pseudogene functions found here to be over-represented among human isolates; this group included some virulence factors (e.g., effector protein YopJ, type IV secretion protein Rhs) (Table S3), suggesting that these virulence functions are more important in bovine than human hosts.

**Figure 3.**
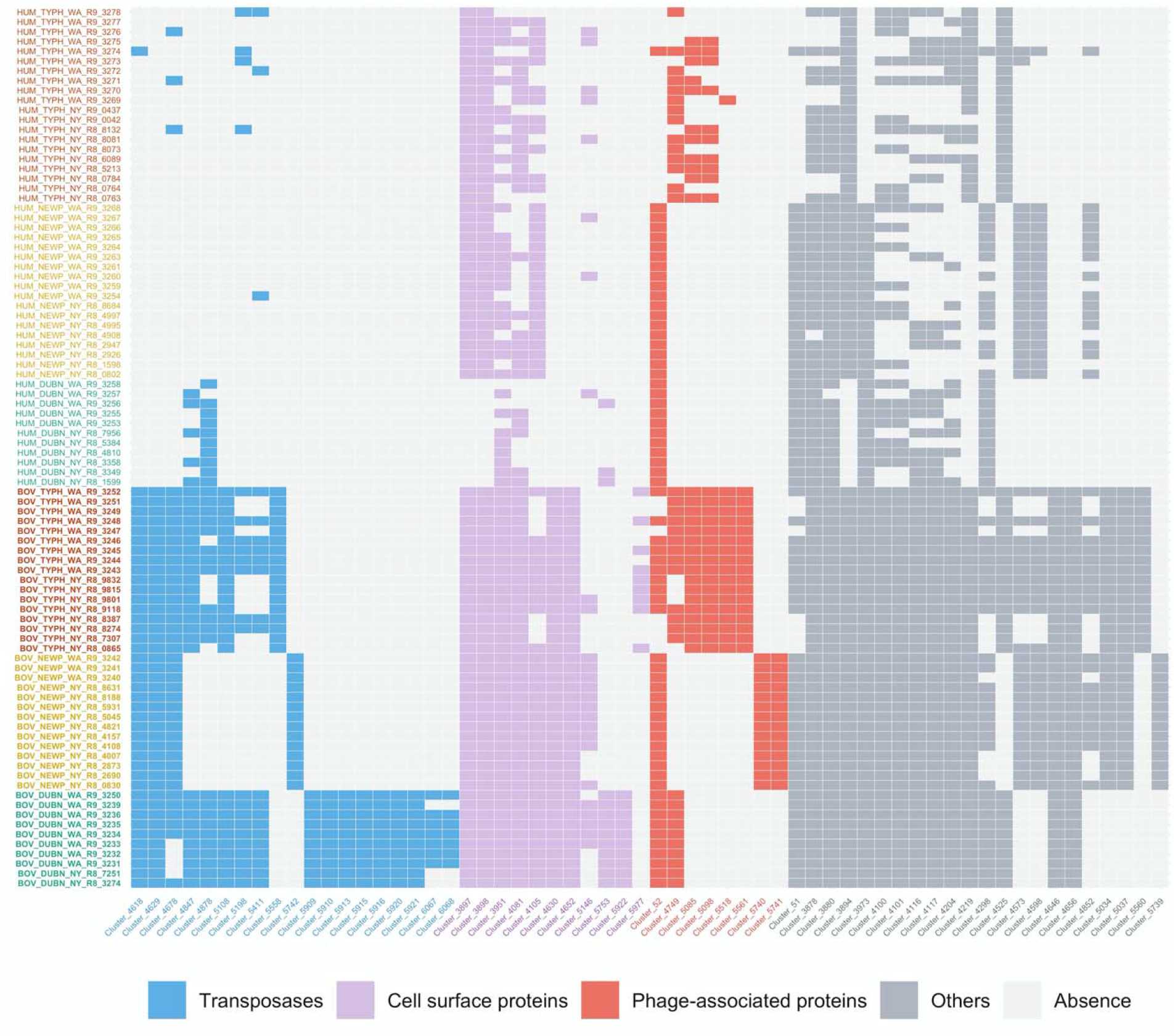
Heatmap of the 60 bovine-associated of AMR *S. enterica* orthologous genes annotated with known functions. Genes encoding transposases, cell surface proteins, and phage-associated proteins are highlighted in blue, purple, and red, respectively; genes encoding other functions are indicated in dark grey. Light grey boxes indicate absence of genes (see key). Isolates (shown on y axis) are color coded by serotype, with green indicating *S.* Dublin, yellow indicating *S.* Newport, and orange indicating *S.* Typhimurium; bovine isolates are highlighted in bold. Details of these 60 bovine-associated genes are shown in Table S3.

Enrichment analysis of GO/EC terms identified 8 terms that were significantly enriched among the 41 bovine isolates representing all three serotypes, including the terms “transposition” (GO:0032196), “double-stranded methylated DNA binding” (GO:0010385), and “hemi-methylated DNA-binding” (GO:0044729) (Figure S1b, Table S4). Within *S.* Dublin, *S.* Newport, and *S.* Typhimurium, a total of 9 (e.g., GO:0050545:endopeptidase inhibitor activity, GO:0046148: pigment biosynthetic process, GO:0032196:transposition), 1 (GO:0032196: transposition), and 1 (GO:0032196: transposition) GO/EC terms were significantly enriched in bovine isolates, respectively (Table S4).

### Bovine isolates exhibited significantly more transposase genes but fewer transposase pseudogenes than human isolates

As a large number of transposases from different IS families were identified as human-associated pseudogene functions (Table S1) and many active transposase genes were enriched among bovine isolates (Figure 3), further analysis was conducted on the comparison of number of pseudogenes as well as orthologous genes (which represent “non-pseudogenes”) annotated as encoding a transposase between bovine and human isolates. Results showed that there were significantly more transposase pseudogenes in human isolates (14, 20, 11, and 14 on average) than bovine isolates (4, 4, 5, and 4 on average) across all serotypes, within *S.* Dublin, *S.* Newport, and *S.* Typhimurium, respectively (Figure 4a, *P* < 0.0001). By contrast, orthologous genes annotated as transposase showed an opposite pattern. There were significantly more genes annotated as transposase among bovine isolates with an average of 19, 29, 13, and 17 than human isolates with an average of 13, 16, 9, and 15 across all serotypes *(P* < 0.0001), within *S.* Dublin *(P* < 0.0001), *S.* Newport *(P* < 0.0001), and *S.* Typhimurium *(P* < 0.05), respectively (Figure 4b). Hence, both pseudogenes and genes annotated as transposases exhibited strong host associations.

**Figure 4.**
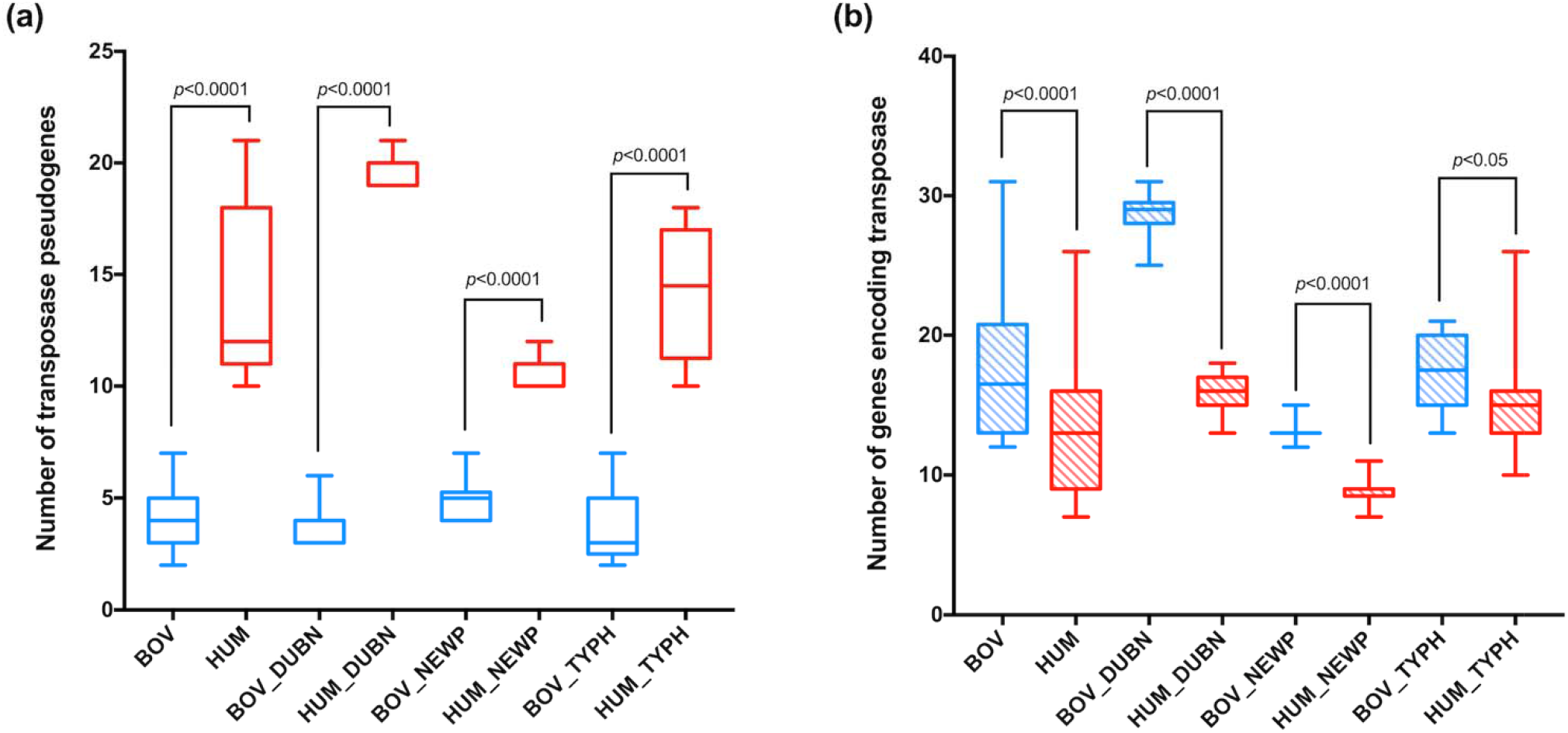
Box and whisker plot of number of (a) pseudogenes and (b) orthologous genes annotated as encoding transposases in AMR *S. enterica* isolates from bovine and human populations. Bovine isolates are indicated by blue boxes; human isolates are indicated by red boxes; *P* values were determined by Mann-Whitney test after FDR correction. Minimum and maximum values are depicted by short horizontal lines above and below the box; the box signifies the upper and lower quartiles, and the median is represented by a short line within the box.

Among all 69 orthologous transposase genes (i.e., genes present in at least one of the 90 isolates studied), 39 showed strong positive correlation with AMR genes, including 16 identified as bovine-associated transposase genes (Phi coefficient r > 0.25, Figure 5a; Table S5). Specific examples of high correlations (Phi coefficients r > 0.8) between transposase genes and AMR genes include (i) one IS91 family transposase gene (Cluster_3939) and *floR* (resistance to phenicols); (ii) one IS6 family transposase genes (Cluster_3940) and *sulII* (resistance to sulfonamides) as well as *strA-B* (resistance to aminoglycosides); (iii) one IS6 family transposase genes (Cluster_5580) and *tetRG* (resistance to tetracyclines), *tetG* (resistance to tetracyclines) as well as *blaCARB* (resistance to beta-lactam); and (iv) one IS1380 family transposase gene (Cluster_4114) and *sulII* (resistance to sulfonamides).

**Figure 5.**
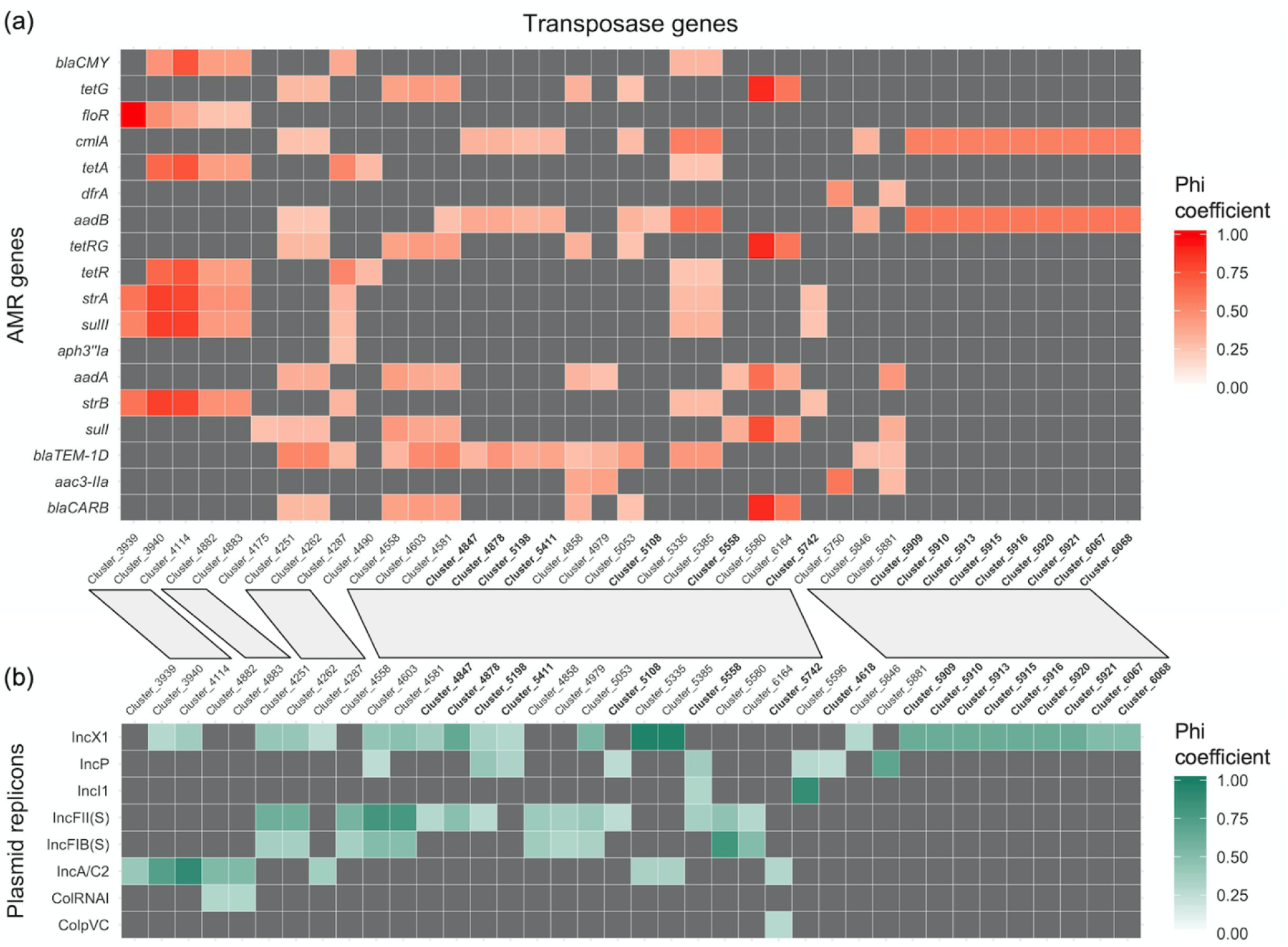
Heatmaps of Phi coefficient between (a) transposase genes and AMR genes, and (b) transposase genes and plasmid replicons. Dark grey boxes indicate that the positive correlation is not strong (Phi coefficient < 0.25). Bovine-associated transposase genes are highlighted in bold. Gray parallelograms between panels a and b link clusters (e.g., Cluster_3939) that showed strong positive correlations with both AMR genes and plasmid replicons.

Phi coefficient analysis was further performed to assess the correlation between transposase genes and plasmid replicons. A total of 38 transposase genes showed strong positive correlation with plasmid replicons, including 17 identified as bovine-associated transposase genes (Phi coefficient r > 0.25, Figure 5b; Table S6). Specific examples of high correlations (Phi coefficients r > 0.8) between transposase genes and plasmid replicons include (i) one IS1380 family transposase gene (Cluster_4114) and IncA/C2; (ii) one IS110 family transposase gene (Cluster_5385) and IncX1; (iii) one IS6 family transposase gene (Cluster_5580) and IncFIB(S); (iv) three transposase genes with no specific family annotated (Cluster_4603, Cluster_5335, and Cluster_5596) and IncFII(S), IncX1, and IncI1, respectively. The majority (36) of these 38 transposase genes positively correlated with plasmid replicons were also positively correlated with AMR genes (Figure 5), suggesting plasmid-mediated transposons potentially carrying AMR genes. Notably, 9 transposase genes showing the same pattern in the correlation with AMR genes (all positively correlated with *cmlA* and *aadB*) were very likely located on plasmid IncX1 (these genes are shown on the right-hand side of Figure 5).

### A larger number of genes and GO terms were associated with human isolates, as compared to bovine isolates

With the same types of analyses described above for identification of bovine-associated genes, a larger number of human-associated genes were found (Figure S1a, Table S2). As compared to 118 bovine-associated genes identified in a combined analysis of all 90 isolates representing the three serotypes, a total of 390 genes were significantly enriched in human isolates across all serotypes (FDR < 0.05, odds ratio > 6.71), including 119 genes present in all 49 human isolates, but absent from all 41 bovine isolates (Figure S1a; see Table S2 for a list of all genes). Annotated functions associated with these 119 genes predominantly represented the category “hypothetical proteins” (99 of the 119 genes); examples of annotated functions in this group of genes included genes encoding an arsenic transporter, a ferredoxin, and a secreted effector protein SteB (Table S2).

Within *S.* Dublin, *S.* Newport, and *S.* Typhimurium, a total of 249, 244, and 244 genes, respectively, were significantly enriched in human isolates (FDR < 0.05, odds ratio > 6.71), including 222, 195, and 188 genes found in all human isolates representing a given serotype, but not found in any bovine isolates with a given serotype (Figure S1a, Table S2). Based on these data, we identified an overall number of 413 human-associated genes, which included the 390 genes identified as human-associated across all serotypes as well additional human-associated genes found within each serotype. These 413 human-associated genes included 278 annotated as “hypothetical proteins” as well as 135 genes with known functions; a large number of genes (n= 42) were annotated as encoding cell surface proteins (e.g., transporters, secretion system proteins), phage-associated proteins (e.g., integrase, phage tail protein), and transcriptional regulators (e.g., LysR, Cro/CI, AraC, LuxR) (Figure 6, Table S7). The annotated functions for 43 human-associated genes matched pseudogene functions found here to be over-represented among bovine isolates. As this group includes genes previously suggested to be involved in host infection (e.g., fimbrial protein FimI, deubiquitinase SseL, heme ABC transporter ATP-binding protein CcmA) (Table S7), our data suggest that these genes may be more important for human than bovine infections.

**Figure 6.**
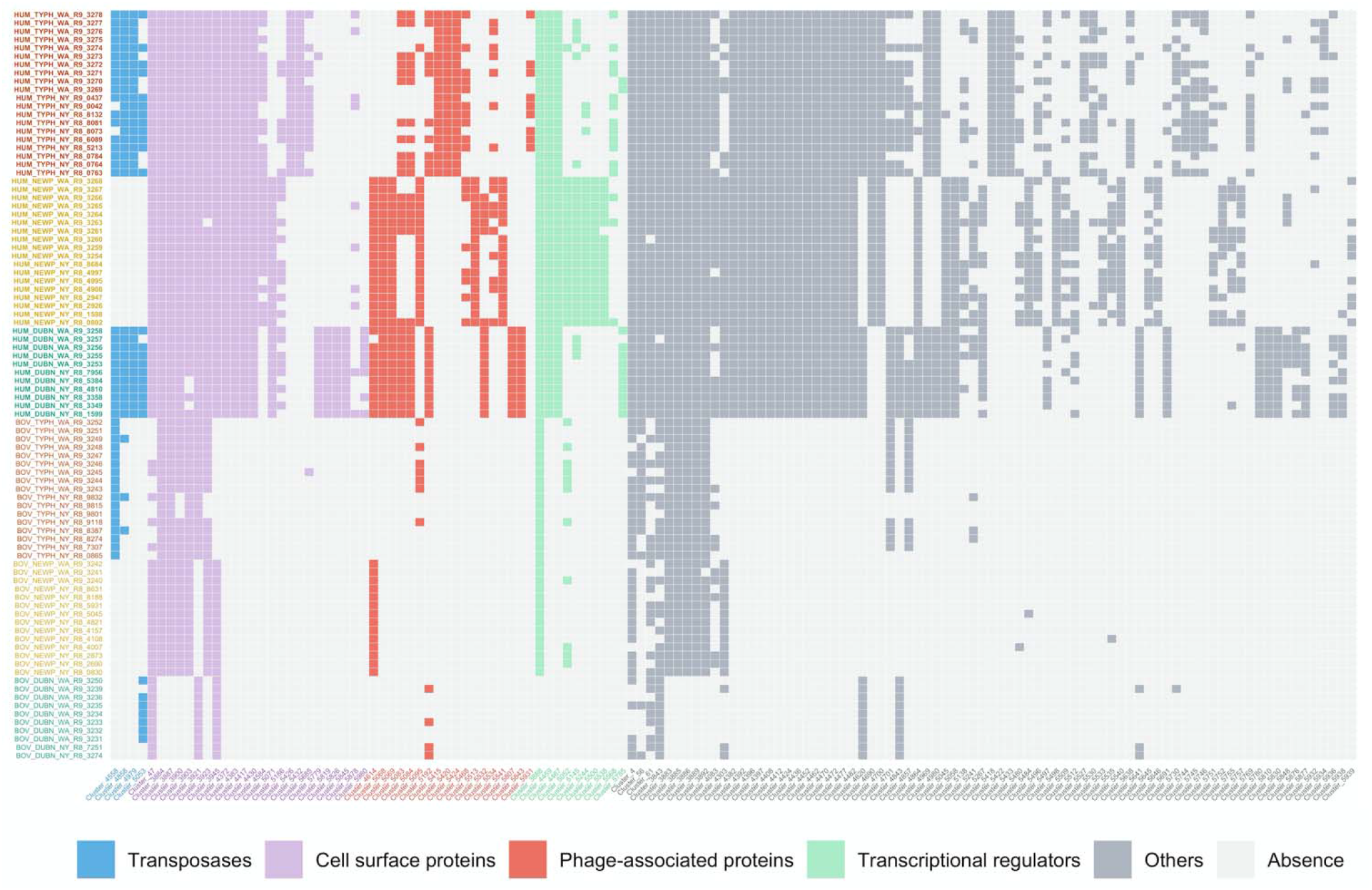
Heatmap of the 135 human-associated AMR *S. enterica* orthologous genes annotated with known functions. Genes encoding transposases, cell surface proteins, phage-associated proteins, and transcriptional factors are highlighted in blue, purple, red, and green, respectively; genes encoding other functions are indicated in dark grey. Light grey boxes indicate absence of genes (see key). Isolates (shown on y axis) are color coded by serotype, with green indicating *S.* Dublin, yellow indicating *S.* Newport, and orange indicating *S.* Typhimurium; human isolates are highlighted in bold. Details of these 135 human-associated genes are shown in Table S7.

At the gene ontology level, more human-associated GO and EC terms were identified compared to bovine-associated ones (Figure S1b, Table S4). A total of 30 GO/EC terms were significantly enriched in human isolates among all three serotypes, including the GO term “provirus excision” (GO:0032359). While two GO terms were exclusively detected in all 49 human isolates (GO:0071287 cellular response to manganese ion; GO:0030026 cellular manganese ion homeostasis), these GO terms only represent a single protein (the small protein MntS, which is classified into both GO terms) (Table S4). Within *S.* Dublin, *S.* Newport, and *S.* Typhimurium, a total of 17, 11, and 11 GO/EC terms were significantly enriched in human isolates, respectively.

### Potentially geographic-and host-associated SNPs were identified within each serotype

No geographic-associated genes or GO/EC terms were identified across all three serotypes and within each serotype. Thus, genetic variants at SNP level were further investigated. A total of 47,913 polymorphic sites were identified among all 3,317 core genes of the 90 AMR *S. enterica* isolates. No SNPs were shown to have a distribution significantly dependent on the geographic origin and host of isolates across all serotypes based on Fisher’s exact tests (FDR > 0.05). Potentially geographic- and host-associated SNPs were subsequently investigated within each serotype.

Within *S.* Dublin, 342 polymorphic sites were identified in 186 Dublin core genes. Fisher’s exact test identified 13 SNPs at which the distribution of nucleotides was significantly dependent on the geographic origin of isolates (FDR < 0.05). The most prevalent nucleotide differed between isolates from NY and WA for each of these 13 SNPs, and 10 of them resulted in nonsynonymous substitutions (Table 2). Five potentially geographic-associated SNPs in *S.* Dublin were located in genes annotated as encoding cell surface proteins such as conjugal transfer protein, porin, and fimbrial protein SefA (Table 2). By contrast, no potential host-associated SNPs were found within *S.* Dublin (FDR > 0.05).

**Table 2.** Annotated functions of genes which contain potentially geographic-associated SNPs within *S.* Dublin and *S.* Newport

| Gene | Function<sup>a</sup> | Codon with nucleotide enriched in NY (AA)<sup>b</sup> | Codon with nucleotide enriched in WA (AA)<sup>b</sup> |
| --- | --- | --- | --- |
| *S. Dublin* |  |  |  |
| Cluster_82 | integral component of membrane<sup>#</sup> | **CAG (Gln)** | **CGG (Arg)** |
| Cluster_208 | integral component of membrane<sup>#</sup> | **AAA(Lys)** | **GAA(Glu)** |
| Cluster_104 | 4-(cytidine 5'-diphospho)-2-C-methyl-D-erythritol kinase | **CTC (Leu)** | **CCC (Pro)** |
| Cluster_4108 | DNA polymerase III subunit epsilon | **CCC(Pro)** | **TCC(Ser)** |
| Cluster_5206 | hypothetical protein | **CCA(Pro)** | **ACA(Thr)** |
| Cluster_5312 | DUF4755 domain-containing protein | **AGT(Ser)** | **GGT(Gly)** |
| Cluster_3703 | ATP-dependent protease | GCC(Ala) | GCT(Ala) |
| Cluster_4136 | restriction endonuclease subunit M | GCA(Ala) | GCC(Ala) |
| Cluster_4288 | RNA-splicing ligase RtcB | **TAT(Tyr)** | **GAT(Asp)** |
| Cluster_4195 | conjugal transfer protein<sup>#</sup> | **GTA(Val)** | **GCA(Ala)** |
| Cluster_195 | cytosol nonspecific dipeptidase | **GCG(Ala)** | **ACG(Thr)** |
| Cluster_476 | porin<sup>#</sup> | GCA(Ala) | GCT(Ala) |
| Cluster_5407 | fimbrial protein SefA<sup>#</sup> | **CAG(Gln)** | **AAG(Lys)** |
| *S. Newport* |  |  |  |
| Cluster_1979 | transporter<sup>#</sup> | **ATC(Ile)** | **GTC(Val)** |
| Cluster_3964 | phage tail protein | **TTC(Phe)** | **TGC(Cys)** |
| Cluster_3624 | acetyl-CoA carboxylase subunit beta | GCC(Ala) | GCG(Ala) |
| Cluster_2245 | penicillin-binding protein activator | CTA(Leu) | CTG(Leu) |
| Cluster_3735 | cytoplasmic protein<sup>#</sup> | **TAC(Tyr)** | **CAC(His)** |
| Cluster_3207 | cardiolipin synthase B<sup>#</sup> | GGT(Gly) | GGC(Gly) |
| Cluster_1005 | AraC family transcriptional regulator | **TGT(Cys)** | **TAT(Tyr)** |
| Cluster_3568 | 2-succinyl-5-enolpyruvyl-6-hydroxy-3- cyclohexene-1-carboxylic-acid synthase | **GTG(Val)** | **GCG(Ala)** |
| Cluster_815 | DUF179 domain-containing protein | **TGC(Cys)** | **CGC(Arg)** |
| Cluster_2311 | E3 ubiquitin--protein ligase | **TGG(Trp)** | **CGG(Arg)** |
| Cluster_3830 | 4-hydroxyphenylacetate permease<sup>#</sup> | **ACA(Thr)** | **ACC(Thr)** |

<sup>a</sup> # indicates that genes were annotated as encoding cell surface proteins.
<sup>b</sup> SNPs resulting in nonsynonymous substitutions (i.e., change in amino acid (AA)) and relevant AA are in bold; A-Adenine, T-Thymine, G-Guanine, C-Cytosine; Ala-
Alanine, Arg-Arginine, Asp-Aspartate, Cys-Cysteine, Gln-Glutamine, Glu-Glutamate, Gly-Glycine, His-Histidine, Ile-Isoleucine, Leu-Leucine, Lys-Lysine, Phe-
Phenylalanine, Pro-Proline, Ser-Serine, Thr-Threonine, Trp-Tryptophan, Tyr-Tyrosine, Val-Valine.

Within *S.* Newport, 480 polymorphic sites were identified in 204 Newport core genes. Fisher’s exact test identified 11 SNPs at which the distribution of nucleotides was significantly dependent on the geographic origin of isolates (FDR < 0.05). The most prevalent nucleotide differed between isolates from NY and WA for each of these 11 SNPs, and 7 of them resulted in nonsynonymous substitutions (Table 2). Among these 11 potentially geographic-associated SNPs in *S.* Newport, 4 SNPs were located in the genes annotated as encoding cell surface proteins (e.g., transporter and 4-hydroxyphenylacetate permease; Table 2). By contrast, no potential host-associated SNPs were found within *S.* Newport as well (FDR > 0.05).

Within *S.* Typhimurium, 2,590 polymorphic sites were identified in 1,425 Typhimurium core genes. While no geographic-associated SNPs were identified within *S.* Typhimurium (FDR > 0.05), 537 SNPs were found to have a distribution of nucleotides significantly dependent on the isolate host (FDR < 0.05). All 537 potentially host-associated SNPs of *S.* Typhimurium had the same most prevalent nucleotide among bovine and human isolates, except for one SNP located in a gene annotated as encoding a hypothetical protein. The majority of those SNPs (77%) were found in 3 genes annotated as encoding hypothetical proteins and 2 genes annotated as encoding transcriptional regulator and NAD(P)-dependent oxidoreductase, respectively. For a total of 523 of the 537 SNPs (97%) a monophyletic group of 8 human isolates (Figure 5; Liao *et al* (2019)) harbored a different nucleotide compared to 12 other human isolates and all 17 bovine isolates, indicating that the significant host association of these SNPs likely represents a single human host-associated Typhimurium lineage.

In summary, 24 SNPs potentially associated with geographic origin were identified within *S.* Dublin and *S.* Newport; 9 SNPs (38%) were located in genes annotated as encoding cell surface proteins (Table 2). The majority of these SNPs (71%) resulted in nonsynonymous substitutions (Table 2), including one located in a gene (annotated as encoding E3 ubiquitin--protein ligase) showing evidence for positive selection as reported by Liao *et al.*, 2019 using the same set of isolates. While a number of potentially host-associated SNPs were identified within AMR *S.* Typhimurium, those were likely just associated with one lineage of AMR *S.* Typhimurium.

### Few closely related isolates were identified between bovine and human and between NY and WA sources based on high-quality core SNPs

Our results provided evidence for associations between genetic variations (e.g., gene presence/absence, SNPs) and isolate sources (e.g. human or bovine), suggesting limited transmission of AMR *Salmonella* between bovine and human populations, and between NY and WA. We thus used high-quality (hq) core SNP data to identify closely related isolates in order to further characterize the frequency of recent transmission events between different sources among the 90 isolates characterized here. Overall, isolates from different geographic origins as well as isolates from different hosts were not very closely related. Only 6, 4, and 0 human isolates for the three serotypes (Dublin, Newport, Typhimurium) showed <10 SNP difference to at least one bovine isolate; 7 human *S.* Typhimurium isolates showed <50 SNP difference to at least one bovine *S.* Typhimurium isolate (see Figure S2a, Figure S2c, Figure S2e for hqSNP distance matrices for comparison between the two hosts for each serotype). For comparison between the two geographic locations, only 2, 1, and 0 isolates from NY for the three serotypes (Dublin, Newport, Typhimurium) showed <10 SNP difference to at least one WA isolate; 2 *S.* Typhimurium isolates from NY showed <50 SNP difference to at least one *S.* Typhimurium isolate from WA (see Figure S2b, S2d, S2f for hqSNP distance matrices for comparison between the two geographic locations for each serotype).

## Discussion

*Salmonella* has a broad host range including humans, other mammals, birds, and reptiles (Eng *et al.*, 2015; Schleker *et al.*, 2012). *Salmonella* can also survive for a long period of time in water (e.g., sewage, freshwater, marine coastal water, and groundwater) (Baudart *et al.*, 2000). This ability of *Salmonella* to survive and multiply in a wide range of ecological niches and to cause disease in diverse hosts is not only practically significant (as many sources can be responsible for a human infection), but also makes this organism a relevant model for studying the population genetics and evolution of environmentally transmitted zoonotic pathogens. In addition, the fact that antimicrobial resistance, including multi-drug resistance, has emerged multiple times in different *Salmonella* serotypes, makes studies on the population structure of different AMR *Salmonella* serotypes important to help inform design of control and prevention strategies and to better understand the evolution and potential for inter-species transmission of AMR zoonotic pathogens with wide host ranges. A number of previous studies have identified genetic markers associated with host adaptation in *S. enterica* (Langridge *et al.*, 2015; Lupolova *et al.*, 2017; Thomson *et al.*, 2008; Yue *et al.*, 2015). For example, Thomson *et al.* (2008) reported that chicken-restricted *S. enterica* serovar Gallinarum isolates showed extensive genome degradation through deletion and pseudogene formation, including the loss of flagella or fimbriae. Lupolova *et al.* (2017) demonstrated that, based on gene content, the source hosts including human, bovine, and swine of *S.* Typhimurium can be accurately predicted using machine learning approach. However, only a few studies, typically focused on non-AMR strains, have explored genetic markers associated with geographic origins of *S. enterica* isolates (Sangal *et al.*, 2010, Zheng *et al.*, 2017). While an initial study has shown that AMR genes of *S. enterica* exhibited a strong association with both host source and geographic origin (Carroll *et al.*, 2017), our study reported here indicates that different evolutionary forces appear to drive AMR *S. enterica* population structure in human and bovine hosts and geographic regions. Specifically, our data not only suggests that transposition plays a more important role in genome evolution and antibiotic resistance transmission of contemporary bovine AMR isolates, as compared to human isolates, but also suggests that association of different virulence-associated genes and pseudogenes with bovine and human AMR *Salmonella* could represent a barrier to inter-host transmission. Importantly, our findings also suggest that specific genetic markers may be able to help predict host or geographic origin of AMR *Salmonella* isolates.

### The host-transposase association suggests that the occurrence time of large-scale transposition and adaptation strategies differ by AMR *S. enterica* populations from bovine and human

Transposase is a DNA-binding enzyme that binds to the end of a transposon and catalyzes the movement of the transposon to another location of the genome via ‘cut-and-paste’ or ‘copy-and-paste’ mechanisms (Rice and Baker, 2001). The significant enrichment of intact genes annotated as encoding transposase among bovine isolates suggests ongoing large-scale transposition and genomic rearrangement events in bovine AMR *S. enterica* populations. By contrast, those events probably occurred earlier in human AMR *S. enterica* populations, since a substantial number of genes annotated as encoding transposase represent pseudogenes in genomes of human AMR *S. enterica* and were not functional anymore. Consistent with our findings, bovine *S. agalactiae* genomes showed a significantly elevated number of transposable elements compared to human *S. agalactiae* genomes (Richards *et al.*, 2011), indicating that such distribution patterns of transposase influenced by host is not restricted to *S. enterica*. This phenomenon might be caused by the different stages of host adaptation/restriction pathogens are experiencing in different hosts. As reviewed by Moran and Plague (2004), when bacteria switch from a free-living to host restriction lifestyle, IS elements in genomes are expected to proliferate during the short-term process of host dependence, with the purpose of reducing genome size by disrupting existing regulatory regions and inactivating genes via transposons (Feschotte, 2008); in the long run, mobile elements are gradually deleted or mutated by forming pseudogenes due to lack of exposure to novel element transposition. As such, facultative intracellular species including *S. enterica* normally harbor dramatically more transposase genes than obligate intracellular bacteria (e.g., *Wigglesworthia glossinidia, Buchnera aphidicola, Chlamydophila caviae*) (Bordenstein and Reznikoff, 2005). Additionally, high numbers of transposase genes inactivated by forming pseudogenes have been observed in the luminous bacterial symbionts of deep-sea ceratiod anglerfishes (Hendry *et al.*, 2018). Based on these mechanisms of genomic changes and previous findings, our data suggests that AMR *S. enterica* in human populations may be at a later stage of host adaptation than those in bovine populations. Adaptation to a human host has also been observed in other host generalist serotypes of *Salmonella*. Klemm *et al.* (2016) detected substantial genome degradation via pseudogene formation and an elevated substitution rate in *S.* Enteritidis which had infected an immunocompromised human patient for 15 years, highlighting the fast within-human evolution of *Salmonella*.

Besides potentially taking advantage of transposons to degrade the genome during a likely early stage of host adaptation, it is also possible that bovine AMR *S. enterica* are maintaining a large number of transposase genes in their genomes due to some beneficial mutations responsible for adaptive phenotypic changes brought by transposons (Casacuberta and González, 2013). One such benefit may be promoting bacterial adaptation to antibiotics used in clinical practice and agriculture (Blot, 1994). Our data showed that more than half of the transposase genes identified in this study showed a significant positive correlation with AMR genes. Congruous with our finding, a variety of transposons carrying antimicrobial resistance genes have been identified in *Salmonella* as summarized in Miriagou *et al.*, (2006). For example, the tetracycline resistance gene *tet*(A), which encodes a membrane-associated efflux protein, has been reported to be linked to transposon Tn1721 (Pezzella *et al.*, 2004). *bla*_TEM-1_ which encodes β-lactamases that mediate resistance to β-lactam antimicrobial agents, has been shown to be carried by Tn3 (Pasquali *et al.*, 2005). The *strA-strB* genes, which confer streptomycin resistance, have been described as part of a particular Tn5393-derivative transposon (Carattoli *et al.*, 2002). The *armA* gene, which encodes aminoglycoside resistance methylase and confers resistance to 4,6-disubstituted deoxystreptamines and fortimicin, has been described as linked to a composite transposon Tn1548 (Galimand *et al.*, 2005). Class 1 and class 2 integrons, which often harbor gene cassettes encoding aminoglycoside modifying enzymes, are commonly associated with various transposons of the Tn3 family (e.g., Tn21, Tn1696, and Tn1412) and Tn7 transposon, respectively (Miriagou *et al.*, 2006). In our study, *tet*(A) was found to be highly correlated with IS1380 family transposase genes; *bla*_TEM-1D_ was highly correlated with IS200/IS605 family transposase, IS630 family transposase, and other two non-specific family annotated transposase genes; *strA-strB* were highly correlated with IS6 family transposase and IS1380 family transposase genes. Selection of antibiotic resistance genes carried by transposons is a strong force in maintaining transposons in populations (Blot, 1994). Thus, keeping a large number of transposase genes could be a strategy employed by bovine AMR *S. enterica* to better adapt to the antibiotics used in cows.

### Identification of host associated genes and pseudogenes encoding presumptive virulence factors suggests adaptation of AMR *S. enterica* to human and bovine hosts

Our analyses identified a number of host-associated genes, including genes found exclusively in isolates from one host (i.e., human or bovine) as well as genes that more frequently presented premature stop codons (hence forming pseudogenes) in isolates from one host. These findings provide initial evidence for adaptation of AMR *S. enterica* to bovine and human populations, at least in the isolate set analyzed here. More specifically, identification of host-association in genes annotated as encoding potential virulence factors supports such adaptation. We specifically identified a number of putative virulence factor-encoding genes that were overrepresented among human AMR *S. enterica* relative to bovine isolates, suggesting not only a specific role of these genes in human infections, but also indicating the possibility of reduced ability to infect human hosts among at least some bovine isolates. For example, the genes encoding the deubiquitinase SseL and fimbrial protein FimI were found to be significantly enriched in human isolates; on the other hand, *sseL* and *fimI* pseudogenes were significantly enriched in bovine isolates. Deubiquitinase SseL, one of the *Salmonella* SPI-2 Type III secretion system effectors, shows deubiquitinating activity during infection of human and muriune cell lines (Rytkönen *et al.*, 2007). Jennings *et al.* (2017) furthermore reported that SseL inhibits accumulation of lipid droplets, prevents autophagic clearance of cytosolic aggregates, and induces late macrophage cell death. *fimI* is required for production of normal type 1 fimbriae and is located in the *fim* gene cluster (Knight and Bouckaert, 2009). It is currently not clear whether FimI constitutes a subunit of type 1 fimbriae or is a protein regulating the assembly of type 1 fimbriae (Knight and Bouckaert, 2009). It has been reported that mannose-sensitive type 1 fimbriae of *S.* Typhimurium promote invasion and appear to play a critical role as an accessory virulence factor (Ernst *et al.*, 1990). As mutations in *fimI* lead to dysfunctional type 1 fimbriae (Valenski *et al.*, 2003), absence of functional FimI (e.g., through formation of pseudogenes, as found here among bovine isolates) may lead to reduced virulence at least in some hosts. We also found that the gene encoding SteB is exclusively found in all human isolates but absent from all bovine isolates in this study. SteB is an effector that requires *Salmonella* SPI-1or SPI-2-encoded T3SS for its translocation (McGhie *et al.*, 2009). SteB has been reported to act as a putative picolinate reductase which is required for efficient mouse spleen colonization in *Salmonella* (McGhie *et al.*, 2009). Identification of putative and well-documented virulence factors encoded by human AMR *S. enterica*, but absent or inactivated in bovine isolates not only indicates that these genes encode host specific virulence factors, but also suggests adaptation of *Salmonella* to human hosts.

In addition, we also identified some virulence genes associated with bovine hosts. For example, the genes annotated as encoding the type IV secretion protein Rhs and the effector protein YopJ were not only found to be significantly enriched in bovine isolates, but *rhs* and *yopJ* pseudogenes were also overrepresented among human isolates. *rhs* genes were first described in *E. coli*, and subsequently found in a wide range of Gram-negative bacteria, including other members of the *Enterobacteriaceae* (e.g., *Salmonella*) as well as *Pseudomonadaceae*. Rhs has been shown to be a mammalian virulence determinant in *Pseudomonas aeruginosa*; Rhs was found to be induced during infection of monocyte/macrophage-like cells, was translocated into these cells, and subsequently caused inflammasome-mediated cell death (Kung *et al.*, 2012). YopJ effectors have been found in a variety of animal pathogens including *Yersinia* spp., *S. enterica*, *Vibrio parahaemolyticus* and *Aeromonas salmonicida* (Ma and Ma, 2016). These effectors target proteins in hosts by acetylating specific serine, threonine, and/or lysine residues, thus influencing the function and/or stability of their target proteins and eventually dampening innate immunity of hosts (Ma and Ma, 2016). Our findings suggest that the type IV secretion protein Rhs and the effector protein YopJ may play a more important role in AMR *S. enterica* during infection of bovine hosts than they do in humans. Consistent with our results, *S.* Typhimurium has been shown to use host-specific bacterial factors (e.g., type III secretion systems, cell surface polysaccharides, cell envelope proteins) to colonize calves and chicks (Morgan *et al.*, 2004).

Furthermore, we identified a number of genes without likely virulence-associated functions that were associated with either bovine or human isolates. While many annotated bovine-associated genes represented transposases, a number of human-associated genes represented other functions. For example, the small protein MntS, which helps to enlarge the manganese pool when manganese is scarce in *E. coli* (Martin *et al.*, 2015), was encoded by a gene exclusively found in all human AMR *S. enterica* isolates but absent from all bovine isolates in this study. *ccmA* was also significantly associated with human isolates, while a *ccmA* pseudogene was overrepresented among bovine isolates; *ccmA* encodes the heme ABC transporter ATP-binding protein CcmA, a putative ATPase essential for TMA/TMAO and nitrite/nitrate as terminal electron acceptors under conditions of reduced oxygen (Batista *et al.*, 2015). Interestingly, all *S.* Pullorum genomes analyzed by Batista *et al.* (2015) also carried a *ccmA* pseudogene; *S.* Pullorum is highly adapted to poultry. In addition, a large proportion of human-associated genes in our study reported here were annotated as encoding transcriptional regulators (e.g., AraC, LuxR, Rha family transcriptional regulator) and phage-associated proteins; Lupolova *et al.* (2016) previously also reported that human *E. coli* O157 isolates showed a higher frequency of phage-associated genes relative to bovine isolates. The specific roles of those host-associated genetic variants in differential adaptation in *S. enterica* remain unknown and further exploration is needed.

In summary, our findings suggest that AMR *S. enterica* serotypes Dublin, Newport, and Typhimurium show evidence of adaptation to human and bovine hosts, including through development of repertoires of host-specific virulence genes. While this raises the intriguing possibility that genetic mechanisms may restrict the transmission of AMR *Salmonella*, at least for the three serotypes studied here, between human and bovine hosts, further work on larger isolate sets, including those from different regions, are needed to confirm this hypothesis. Importantly, however, our findings are consistent with other studies which show the development of host-specific subgroups within closely related pathogen taxa. For example, different alleles of *fimH* encoding type 1 fimbrial adhesin showed different binding preference to human cells and bovine cells (Yue *et al.*, 2015). Evolution of host specificity has also been well established in a number of other bacterial pathogens, as reviewed by Bäumler and Fang (2013).

### The geographic-associated SNPs highlight the role of geographic origin in influencing population structure of AMR *S. enterica* and evolution of cell surface proteins

Potential geographic-associated SNPs were found within AMR *S.* Dublin and AMR *S.* Newport. *S.* Newport has been reported to display a geographic structure (Sangal *et al.*, 2010). Specifically, Lineage I strains were more frequently isolated from Europe, while Lineages II and III strains were more likely isolated from North America. Such geographic association of lineages was suggested to be caused by differences in prophage regions, pathogenicity islands and fimbrial operons among strains from different geographic locations (Zheng *et al.*, 2017). AMR *S.* Newport analyzed here belong to Lineage IIC as previously reported by Liao *et al.* (2019). Our findings suggest that within this lineage, *S.* Newport also has a geographic structure at the SNP level.

A large proportion of geographic-associated SNPs found in *S.* Dublin and *S.* Newport (38%) were located in genes annotated as encoding cell surface proteins (e.g., cardiolipin synthase B, porin, permease, transporter, fimbrial protein SefA), higher than the proportion of cell surface proteins among the total amount of proteins in eukaryotic and prokaryotic cells in general (20%-30%). Interestingly, cardiolipin synthase B, which catalyzes the synthesis of cardiolipin in bacteria, has been found to be important in modulating the physical properties of membranes in response to environmental stress such as osmotic stress (Mileykovskaya and Dowhan, 2009). Mutation of cardiolipin synthase has been shown to be associated with daptomycin resistance in enterococci. Davlieva *et al.* (2013) reported that when challenging *Enterococcus faecium* and *Enterococcus faecalis* with daptomycin, the pool of phosphatidylglycerol was substantially reduced. However, mutations observed in gene encoding cardiolipin synthase were able to compensate for the decreased phosphatidylglycerol and lead to a restoration of the cardiolipin synthase pool. In addition, Peleg *et al.*, 2012 found genomic evidence that point mutations in gene coding for cardiolipin synthase were responsible for developing reduced susceptibility to daptomycin in *Staphylococcus aureus*. Daptomycin has been a widely used antibiotic in the treatment of complicated skin and skin structure infections in humans (Steenbergen *et al.*, 2009). While highly speculative, the geographically associated SNPs identified in genes encoding cardiolipin synthase B for *S.* Newport could suggest possible follow-up studies on use of daptomycin in NY and WA.

Another interesting gene, annotated as encoding an exported protein, E3 ubiquitin--protein ligase, contained a geographic-associated SNP resulting in a nonsynonymous substitution, and this gene showed evidence of positive selection in a previous study using this same set of isolates (Liao *et al.*, 2019). Genes encoding E3 ubiquitin--protein ligase are present in several species of pathogenic bacteria, including *Salmonella* (Quezada *et al.*, 2009), and this protein has been reported to be used as a virulence factor (Maculins *et al.*, 2016). Multiple studies have shown that E3 ubiquitin ligase was used by *Salmonella* to hijack host cells and subvert its ubiquitination pathway (Maculins *et al.*, 2016; Quezada *et al.*, 2009). Ubiquitination systems function in a wide range of cellular processes (e.g., cell cycle and division, immune response and inflammation) in eukaryotic organisms, and can lead to cancer and neurodegenerative disorder when ubiquitination is defective (Finley and Chau, 1991). Thus, expressing and releasing of E3 ubiquitin ligase by *Salmonella* could promote bacterial survival and pathogenicity in the host. A geographically associated SNP found in a gene encoding E3 ubiquitin--protein ligase might be a consequence of positive selection triggered by specific ecological niches associated with NY or WA.

Overall, our findings suggest that, in addition to host, geographic origin is also a critical factor contributing to the population structure, antimicrobial resistance, and pathogenicity of AMR *S. enterica*, likely due to distinct ecological niches (e.g., represented by climate, dairy operation management, antibiotics use) associated with different geographic origins.

### AMR *S. enterica* population structure is likely driven by different evolutionary forces and a low potential of recent transmission among hosts and geographic origins

It has been demonstrated that ecological niches can change population structure and affect the evolution of bacteria (Vos, 2011; Li *et al.*, 2019). A recent study (Behringer *et al.*, 2018) on the long-term evolution of *E. coli* cultures proposed that the evolution of genetic differentiation resulting in subpopulation structure was facilitated by spatial differentiation followed by metabolic differentiation. To cope with different ecological niches, adaptive evolutionary processes mediated by a mixture of mechanisms - mainly gene flow, mutation, natural selection, and genetic drift – will occur in bacteria, resulting in gene gain/loss and functional divergence of existing genes (Toft and Andersson, 2010; Li *et al.*, 2019; Slatkin, 1987; Strachan *et al.*, 2015). Our data provides evidence in strong association of gene gain/loss with host as well as evidence for SNP level divergence of core genes associated with geographic origin (in at least some AMR *S. enterica* serotypes), suggesting different mechanisms are underlying adaptation of AMR *S. enterica* to host and to geographic location. For example, the strong association between bovine-associated transposases and plasmids (e.g., IncX2 and IncFII(S)) observed in this study indicates that some transposons in bovine AMR *S. enterica* were likely obtained via horizontal gene transfer. Consistent with our finding, transposons have been frequently detected on conjugative or mobilizable plasmids in various *Salmonella* obtained from different sources (Miriagou *et al.*, 2006). Most of human-associated pseudogenes are the result of frameshifts and/or premature stop codons caused point mutations, which are common mechanisms in the formation of pseudogenes in bacterial genomes (Lerat and Ochman, 2005). While geographically associated SNPs were not exclusive under positive selection, nonsynonymous SNPs in genes encoding cell surface proteins are potential targets of positive selection, as cell surface proteins have been found to be a major group driven by positive selection in bacteria (Chen *et al.*, 2006; Petersen *et al.*, 2007; Wachter and Hill, 2016). By contrast, synonymous geographic-associated SNPs may be a consequence of genetic drift, as genetic drift is the driving force for neutral mutations (Bromham and Penny, 2003). Overall, our findings suggest that gene flow and mutation are playing important but not exclusive roles in shaping host-associated population structure of AMR *S. enterica*, while natural selection and genetic drift are essential, but not exclusive, in shaping geographically associated population structure of AMR *S. enterica*.

In addition to different evolutionary forces influencing the population structure of AMR *S. enterica*, our data together with previously reported associations between AMR genes and isolate source (Carroll *et al.*, 2017) also suggests a low potential for recent transmission of AMR populations between bovine and human populations and between NY and WA. Transmission of bacteria can promote genetic exchanges, so a more homogenized gene pool and closely related individuals are expected (Slatkin, 1987). In this study, AMR *S. enterica* showed high degree of host-genome and geographic-genome associations at gene, GO terms, pseudogene, and SNP levels, and absence of a large number of closely related isolates between bovine and human populations and between NY and WA. Similarly, Mather *et al.* (2013) demonstrated that *S.* Typhimurium DT104 and its antimicrobial resistance genes were maintained largely separately within animal and human populations in Scotland, with little transmission between animals and humans and vice versa. Overall, these data suggest that transmission of *Salmonella* between bovine and human hots may be less frequent than sometimes assumed, even though there have been cases of AMR *S.* Dublin (Harvey *et al.*, 2017), *S.* Newport (Centers for Disease Control and Prevention, 2002; Plumb *et al.*, 2019), *S.* Typhimurium (McLaughlin *et al.*, 2006; Olsen *et al.*, 2004) outbreaks in humans linked to raw milk, raw beef or cheese. Our findings are also consistent with some studies that indicated geographical association with different *Salmonella* clonal groups; for example, Kovac *et al.* (2017) reported that *S.* Cerro found in Texas and NY represent largely distinct clonal groups. As NY and WA are about 3,500 km apart, it is plausible that such a distance could generate a barrier for *Salmonella* transmission, since microbial dispersal is typically a passive process, which does not likely result in long-distance dispersal events (Nemergut *et al.*, 2013). While contemporary complex food distribution systems may reduce the effectiveness of distance as a barrier for transmission, our data suggest that geographical distance may still represent a barrier to long-distance *Salmonella* dispersal.

## Conclusion

The population structure of antimicrobial resistant bacterial pathogens which differs in host and geographic origin could reveal key elements in understanding their evolution in adaptation, pathogenesis, and antimicrobial resistance. Collectively, our study identified a number of genetic variants in AMR *S. enterica* associated with host and geographic origin using comparative genomics. It highlights a vital role of host in shaping genome arrangement and promoting antimicrobial resistance via potentially plasmid-mediated transposons and in developing host adaption via preferred virulence factors. It also highlights a critical role of geographic origin in driving the evolution of cell surface proteins and antimicrobial resistance potentially mediated by positive selection and genetic drift. Even though the advancement of sequencing techniques has allowed for rapid progress in identifying niche-associated genetic variants, our understanding of the evolutionary mechanisms underlying the generation, fixation and function of those variants in populations is still limited. Our study provides a large number of candidate genes, such as secreted effector protein SteB, fimbrial protein FimI, E3 ubiquitin--protein ligase, for future investigations to better understand the adaptation and pathogenesis of AMR *S. enterica*. As AMR *S. enterica* represent a substantial public health concern, focusing on AMR populations allowed us to identify specific genetic markers that could be potentially used in source tracking (e.g., host and geographic origins) of human disease cases and contamination events caused by AMR *S. enterica*. Our data also suggest a low potential for inter-host and long distance-geographic transmission events of AMR *Salmonella*. Despite the benefits of focusing on AMR populations, this study is limited by a relatively small dataset. To fully understand the genetic basis of environmental adaptation and differential virulence associated with host and geographic origin in *S. enterica*, it is necessary for future studies to include non-AMR populations as well.

## Author Contributions

JL and MW conceived and designed the study. JL wrote the manuscript with input from MW, RHO and LC. JL performed the computational analyses and statistical analyses.

## Acknowledgement

This work was partially supported by the USDA National Institute of Food and Agriculture Hatch project NYC-143436. Any opinions, findings, conclusions, or recommendations expressed in this publication are those of the author(s) and do not necessarily reflect the view of the National Institute of Food and Agriculture (NIFA) or the United States Department of Agriculture (USDA). We are grateful to Dr. Rachel A. Cheng for her assistance in interpreting functions of host-associated genes.

