## Supplementary_File_2 for "Comparative genomics reveals different population structures associated with host and geographic origin in antimicrobial-resistant *Salmonella enterica*"

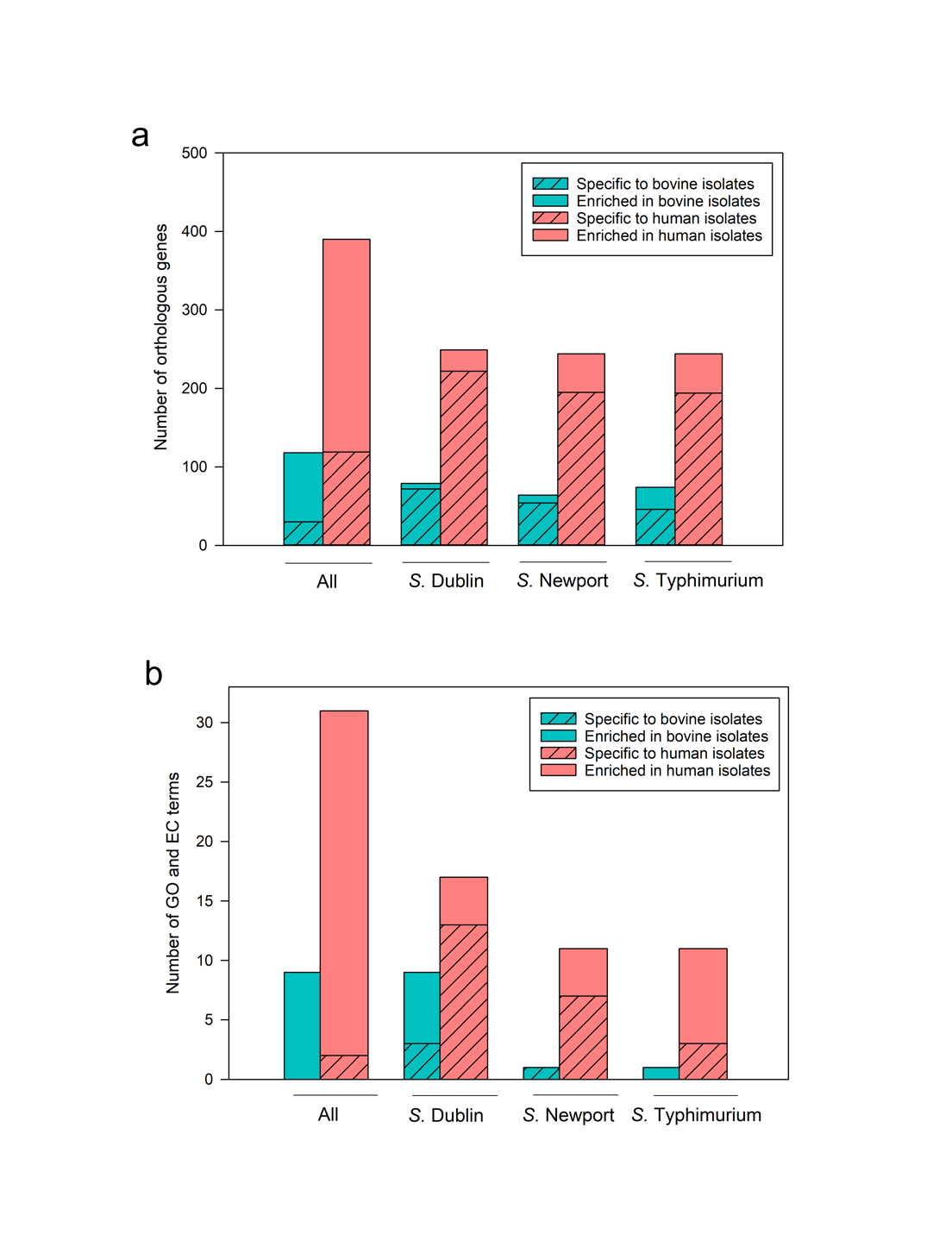

**Figure S1.** Enrichment of (a) orthologous genes and (b) gene ontology (GO) and enzyme commission (EC) terms in bovine and human isolates among all three serotypes and each serotype. Orthologous genes and GO/EC terms significantly enriched in one host type were determined when FDR < 0.05 and odds ratio > 6.71. Bovine-associated orthologous genes and GO/EC terms were indicated by blue bar; human-associated orthologous genes and GO/EC terms were indicated by red bar.

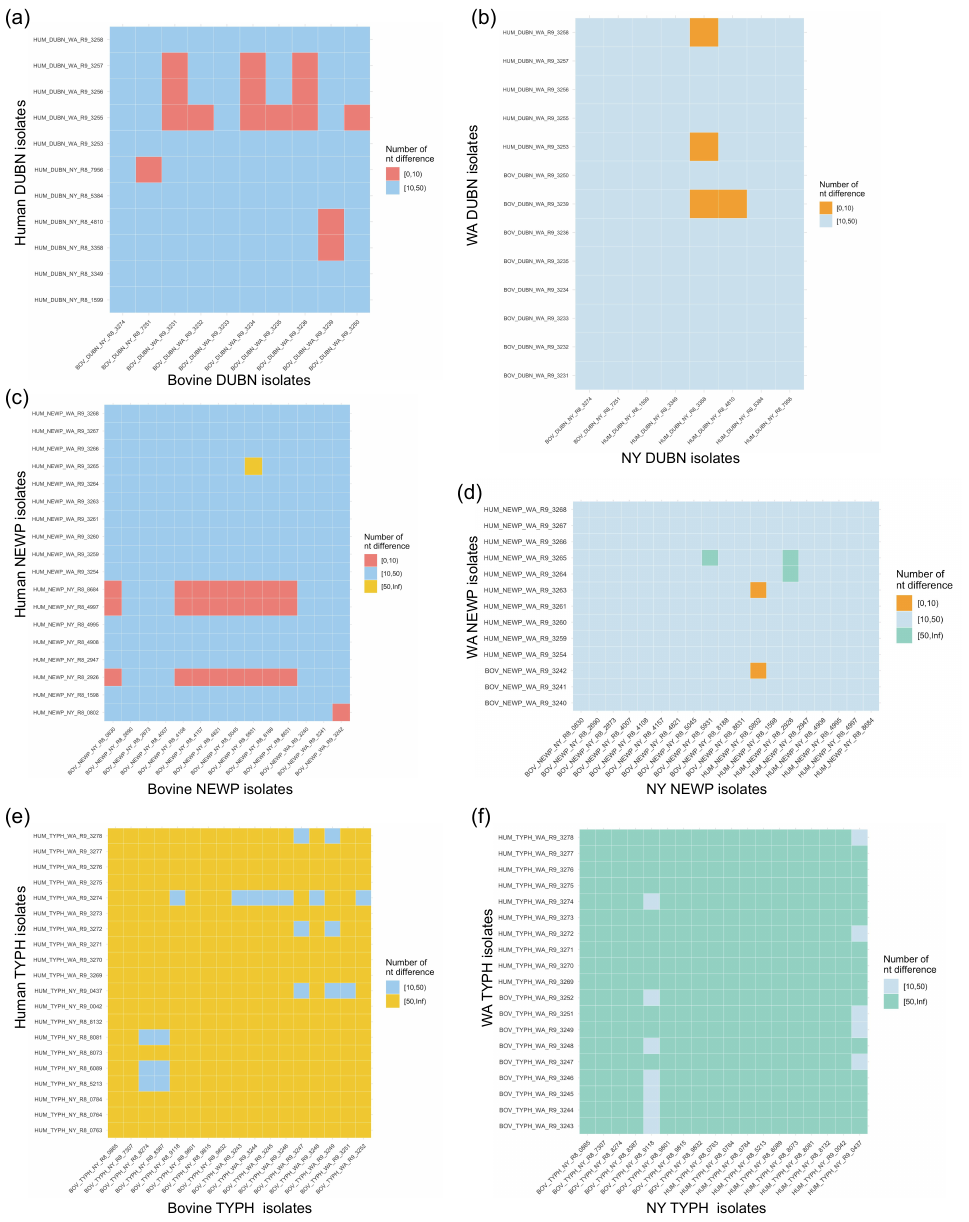

**Figure S2.** Heatmaps of nucleotide difference between (a) bovine and human *S.* Dublin (DUBN) isolates, between (b) NY and WA *S.* Dublin isolates, between (c) bovine and human *S.* Newport (NEWP) isolates, between (d) NY and WA *S.* Newport isolates, between (e) bovine and human *S.* Typhimurium (TYPH) isolates, and between (f) NY and WA *S.* Typhimurium isolates. Nucleotide difference ranging from 0 to 10 is indicated by red cells and orange cells for comparison between hosts and between geographic locations, respectively. Nucleotide difference ranging from 10 to 50 is indicated by dark blue cells and light blue cells for comparison between hosts and between geographic locations, respectively. Nucleotide difference larger than 50 is indicated by yellow cells and green cells for comparison between hosts and between geographic locations, respectively.
